# From larval stage to adulthood? Habitat type-specific chemical fingerprints of fire salamander larvae might give first indication for adult mate preference

**DOI:** 10.1101/2025.01.07.631682

**Authors:** Laura Schulte, Jeanne Friedrichs, Caroline Müller, Barbara A. Caspers

## Abstract

Assortative mating can play a major role in the divergence of a population with gene flow. In a fire salamander population near Bonn, Germany, the larvae differ genetically according to the habitat type they are found in, i.e. ponds or streams. Neither sound nor vision are a source for assortative mating in fire salamanders, but it is known that females can discriminate between sexes and habitat types of the males based on chemical cues. The adults share the same terrestrial habitat which leads to the question whether there is a habitat type-specific chemical fingerprint already existing in the larvae that may be used in the adult stage as the base for assortative mating. We took water samples and captured larvae at two ponds and two streams and used polydimethylsiloxane and TD-GC-MS to collect and analyze samples of the environmental background as well as from the surface of the larvae. We found a significantly different composition of chemical features in the environmental back-ground samples. Additionally, we found the larvae to carry habitat type-specific chemical features. These finding allow to speculate that the features on the larval surface may be used

## Introduction

When populations or subpopulations are exposed to different environmental conditions, divergent selection pressures can eventually lead to reproductive isolation, either directly or indirectly, contributing to the process of speciation. Such speciation can occur in allopatry or sympatry (Schluter, 2001). The divergence of a population that shares the same habitat can also be mediated through selective mating (Bolnick and Fitzpatrick, 2007; Higashi et al., 1999). In this context, assortative mating, i.e. the sexual selection after a specific trait, plays a major role in the divergence of populations with gene flow (Acord et al., 2013; Lewontin et al., 1968; Sachdeva and Barton, 2017). Assortative mating can be based on specific traits that the pair shares (Green, 2019), such as size (Taborsky et al., 2009), morph (Wilson et al., 2004), body condition (Gould and Valdez, 2021) or courtship behaviour (Meguro et al., 2016), but it can also be based on chemical signals (Coombes et al., 2018; Liberles, 2014; Peterson et al., 2007; Caspers & Steinfartz, 2011; Woodley and Staub, 2021). In insects for example, chemical signals, often expressed through cuticular hydrocarbon (CHC) profiles, play a major role in mate choice and can serve as the basis for assortative mating (Ferveur, 2005; Peterson et al., 2007; Singer, 1998). Chrysochus beetles for instance use CHC profiles for male mate choice (Peterson et al., 2007). Likewise, male salamanders of the Western redback salamander (Plethodon vehiculum) and Dunn’s salamander (P. dunni) can discriminate gravid from non-gravid females based on chemical cues (Marco et al., 1998). However, knowledge about the source or what influences the chemical cues is often missing (Jaeger and Gergits, 1979).

Assortative mating can also be mediated through habitat choice (Takahashi, 2004) and can lead to divergence and reproductive isolation. In a cichlid fish system, for instance, differences in courtship behaviour and morphological traits led to assortative mating which then in turn led most likely to sympatric speciation (Barluenga et al., 2006). Additionally, in Midas cichlids (Amphilophus xiloaensis and A. sagittae) assortative mating is based on different colour morphs that share the same habitat (Elmer et al., 2009). Caillaud and Via (2000) demonstrated that pea aphids (Acyrthosiphon pisum) belonging to two different genotypes using different host plants can distinguish between their host plant and a non-host plant. Individuals of each genotype abandon the respective non-host plant while showing assortative mating with conspecifics on their host plant leading to ecological speciation.

In a fire salamander population (Salamandra salamandra) from the Kottenforst near Bonn, Germany, ecological speciation towards the larval habitat type can be observed. The fire salamander has a biphasic lifecycle like most amphibians: The larvae are deposited into water bodies and after metamorphosis the adults are terrestrial except the females that return to water bodies for larval deposition (Thiesmeier, 2004). While fire salamanders typically use first order streams for larval deposition, in the Kottenforst population females also use ponds. Population genetic analysis revealed the existence of two genetic clusters that correlate with the larval habitat type pond or stream (Steinfartz et al. 2007; Hendrix et al. 2017). Interestingly, almost all larvae found in one of the habitat types match genetically the respective habitat-specific genotype (Hendrix et al., 2017; Steinfartz et al., 2007), indicating either assortative mating or selection against larvae of the non-matching genotype.

While distinct habitat types exist for the larvae, adults share the same habitat, independent of the genotype and their larval habitat type. They live in a broadleaf forest that holds several ponds and streams in vicinity to each other. The mating in the fire salamander is mostly terrestrial and not connected to the natal habitat anymore (Caspers et al., 2009; Thiesmeier, 2004, but see Dehling, 2024). Adults are mainly active during the night (Thiesmeier, 2004) and neither sound nor vision play a big role in the communication with conspecifics (Kemp, 2021; Woodley, 2014). Chemicals, however, are very important for intraspecific communication in this species (Kemp, 2021; Caspers & Steinfartz, 2011). Preferences tests, for example, showed that fire salamanders can discriminate the sex of an individual based on chemical cues (Caspers and Steinfartz, 2011), and females show a preference for males of the same habitat type, irrespective of genetic distance/proximity (Caspers et al., 2009), indicating that chemical cues might also allow females to discriminate the habitat type of a potential mate. In other salamander species, females transmit pheromones across their skin during direct contact (Woodley and Staub, 2021). However, it is currently unknown whether such a mechanism is present in fire salamanders. Adults only differ in their habitat of origin, i.e. the larval habitat type, and thus environmental conditions in the adult habitat cannot influence assortative mating. This leads to the question whether the larval habitat type might shape a chemical fingerprint, which allows for habitat type-specific mating once being adult. A pre-requisite for this is that larvae already have a habitat type-specific chemical fingerprint, which is currently unknown.

With this study, we aimed to investigate potential differences in the chemical fingerprints of water bodies and of larval surfaces from the two habitat types. We hypothesised that larvae of the two habitat types, i.e. ponds and streams, differ in their chemical fingerprints. Potential differences in chemical fingerprints of larvae might contribute to intraspecific communication among life stages and might serve as the base for assortative mating in adults that has the potential to lead to ecological speciation.

## Methods

### Sampling in the field

Between 13^th^ – 15^th^ of March 2023 we collected samples from two habitat types, i.e. two pond (P1, P2) and two stream locations (S1, S2) in the Kottenforst, Bonn (Germany)(figure 1). On each location, we collected samples from twelve fire salamander larvae of similar size (2.8 – 4 cm). Each larva was placed in a glass Petri dish that had been cleaned with acetone beforehand and heated up to 200°C to sterilise it. The whole body of the larva was rinsed from the right and the left side of the body starting from the top with distilled water (except for the head and gills) to ensure that we sample only larval-specific features and not features from the water. The chemicals on the surface of each larva were collected by using tubes of absorbent polydimethylsiloxane (PMDS; length 5 mm; diameter: inside 1 mm, outside 1.8 mm; Carl Roth, Karlsruhe, Germany). These PDMS tubes had been cleaned and conditioned beforehand by stirring them for 8 h at 80 °C in an acetonitrile:methanol mixture (4:1, v:v, LC-MS grade, Fisher Scientific, Loughborough, UK), then storing them in fresh solvent for 16 h at room temperature and subsequently heating them at 230 °C for 30 min with a conditioning program at a helium flow of 60 mL min^-1^, following the protocol as described in Kallenbach et al. (2015). The tubes were stored at room temperature in capped glass vials (Macherey-Nagel GmbH & Co. KG., Düren, Germany) sealed with polytetrafluoroethylene (PTFE; Carl Roth, Karlsruhe, Germany) tape until use. In the field, two PDMS tubes were placed on a stainless-steel wire and gently rubbed ten times on each side of the larva. PDMS tubes were only touched with cleaned forceps and the experimenters wore nitrile gloves. Afterwards, tubes were placed again in glass vials closed with caps and sealed with PTFE tape. Additionally, we measured the total length of each larva. Immediately afterwards, each larva was placed back into the water. Then, all the equipment used was cleaned with ethanol (70%).

**Figure 1:**
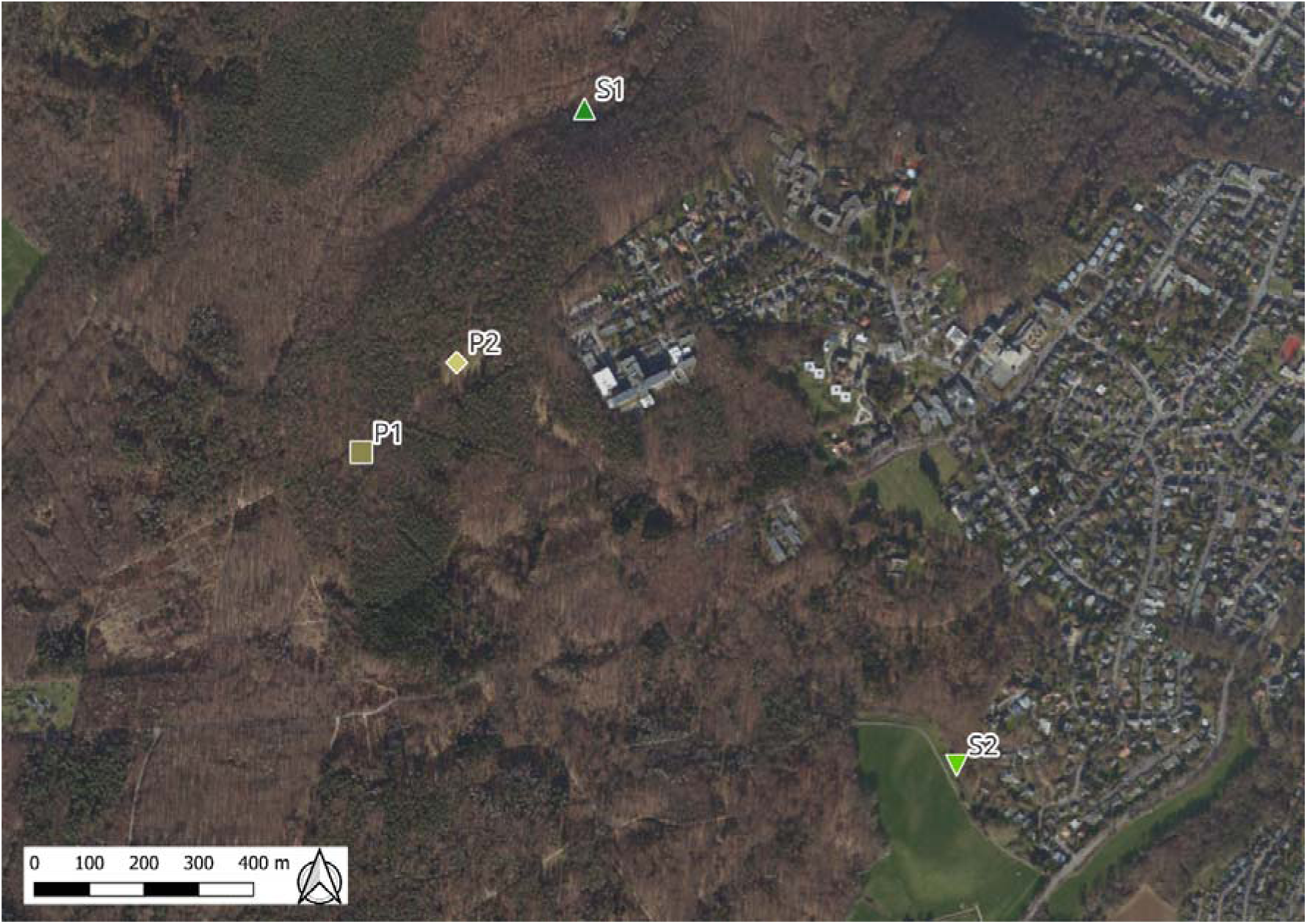
The two locations for the pond habitat type P1 and P2 (brown squares), and the two locations for the stream habitat type S1 and S2 (green triangles), are located in the Kottenforst, Bonn (Germany).

As “handling” samples (controls), we placed three tubes on a wire and dipped those into distilled water, rubbed them ten times over the cleaned glass petri dish and our gloves (one replicate per location). As “environmental background” samples, we placed five tubes on a wire and hold them into the water of each location for 30 seconds (one replicate per location). All samples were immediately stored in the field at -10 °C in a portable freezer and transferred into -20 °C after arriving at the laboratory.

In addition, we also collected abiotic factors (pH, water temperature and oxygen level) at each location on the day of sampling and repeated this every week until mid may, 2023. All measurements were taken using a HANNA HI98196 pH/ORP & dissolved oxygen measurement probe by sampling the parameters at the littoral zone of each water body.

### Analysis of chemical compounds

Surface chemicals collected on the PDMS tubes were analysed in electron impact ionisation mode using thermodesorption gas chromatography mass spectrometry (TD 30–GC 2010 Plus – MS QP2020, Shimadzu, Kyoto, Japan) with a VF-5 MS column (30 m x 0.25 mm ID x 0.25 µm, 10 m guard column, Varian, Agilent Technologies, Santa Clara, California, USA) and helium as carrier gas. PDMS tubes were desorbed at 250 °C under a flow rate of 60 mL min^−1^ and compounds cryo-trapped on Tenax® for 8 min at −20 °C. Compounds were re-desorbed at 250 °C for 3 min, then transferred in a 1:1 split mode, with a transfer line temperature of 250 °C and a column flow rate of 1.72 mL min^−1^ to the GC. The temperature of the GC oven was set to 40 °C for 3 min, increased to 100 °C at 6 °C min^-1^, then to 200 °C at 4 °C min^-1^ and finally to 280 °C in steps of 20 °C min^-1^, holding the final temperature for 5 min. Line spectra (m/z of 30–400) were obtained in quadrupole MS mode at 70 eV with a scan event time of 0.04 sec. The ion source and interface temperature were set to 230 °C and 250 °C, respectively. The PDMS tubes of the handling and environmental background samples were measured individually per location. Additionally, an alkane mix standard (C7-C40, Sigma-Aldrich, St. Louis, USA) and a mix of five standards were measured twice.

Pre-processing of the data was done using the GCMS Postrun Analysis software (GCMSsolution version 4.45, Shimadzu, Kyoto, Japan) by converting the data from .qgd into .CDF file format, and the alignment of peaks was done in R (version 4.3.2, R Core Team 2023) using the packages Bioconducter (BiocManager, version = “3.18”) and metaMS (Wehrens et al., 2014). Features were aligned using the runGC method in the metaMS package with the following adjusted settings in metaMSSettings: method=”matched filter”, step = 0.5, steps = 2, mzdiff = 0.5, fwhm = 3, snthresh = 11, max = 20, CAMERA = list(perfwhm = 10). The metaSettings of alignment “betweenSamples” were min.class.fraction = .01, min.class.size = 1, timeComparison = “rt”, rtdiff = .05, RIdiff = 2 and simthresh = .90. Peak areas of the different features were used as intensities. It has to be noted that this is a rough approximation, as most of the features are unknown and therefore response factors could not be evaluated.

For further data processing, the mean intensity of the features that occurred in the handling samples collected at each location (3 tubes per location) was subtracted from the feature intensities within each salamander sample. Likewise, the mean feature intensities of the handling samples were subtracted from each environmental background sample per location. Additionally, siloxane peaks and other obvious contaminants were removed from the data-set. The remaining feature intensities were then divided by the length of the respective sala-mander larva representing the features of the salamander and the environmental background (see supplements).

### Statistical analysis

To investigate differences in the abiotic parameters (pH, water temperature and oxygen) between the two habitat types, we used t-test or Mann-Whitney-U test depending on the distribution of the data in R Studio for the data collected over the monitoring period of 10 weeks (R version 4.3.2; 2023-10-31 ucrt). We used t-test and one-way anova to test for differences between the size of the larvae per habitat type and location, respectively. We found that the size of the larvae did not differ between the habitat types (t-test: t=-1.5; p=0.141) but per location, with larvae from stream S1 being significantly larger than larvae from S2 and P2 (p=0.002 and p=0.003, respectively), but not different from P1 (p=0.07).

To analyse differences in the feature diversity and chemical fingerprints, we used the software Primer (version 7.0.23). First, we investigated the α-diversity for the features in the environmental background samples as well as in the larval surface samples by calculating the Shannon Index using the “DIVERSE” function in primer with the following formula:

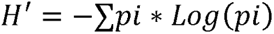

We tested for differences in Shannon Index of the environmental background samples by using the Mann-Whitney-U test and the t-test to test for differences in the Shannon Index in the larval surface samples. After that, we tested for potential habitat type and location differences in the composition of features. We first analysed only the environmental background samples. To do this, we standardised the data and transformed it to presence/absence. Based on a Bray-Curtis similarity matrix we performed an ANOSIM per factor (habitat type/location) as we found no significant difference in the dispersion of the data (Permdisp: habitat type p=0.3, location=0.146).

To analyse the presence/absence of the features found on larvae from ponds and streams, we plotted a venn plot using the packages VennDiagram (Chen and Boutros, 2011) and ggvenn in R (Yan, 2023). Moreover, we tested for potential differences per habitat type and location for the chemical composition of the larval surface samples (presence/absence data) by processing the data as describe above and performing another ANOSIM per factor as we again found no differences in the dispersion (Permdisp: habitat type p=0.343, location p=0.631).

## Results

When comparing the abiotic factors per habitat type, we found ponds to have a significantly lower pH value (Mann-Whitney-U test: p<0.001), but we found no difference in the water temperature between the two habitat types (Mann-Whitney-U test: p=0.144).

In the environmental background samples, we found in total 310 features. We found no difference in the Shannon diversity between the chemical fingerprints of the two habitat types (Mann-Whitney-U test: n=10 per habitat type, p=0.081, Supplements figure 5). In contrast, for the chemical composition of these samples, we found a significant difference between the habitat types (one-way ANOSIM: R=0.177, p=0.004). Likewise, we also found a significant difference for the location (one-way ANOSIM: R=0.279, p=0.001, figure 2).

**Figure 2:**
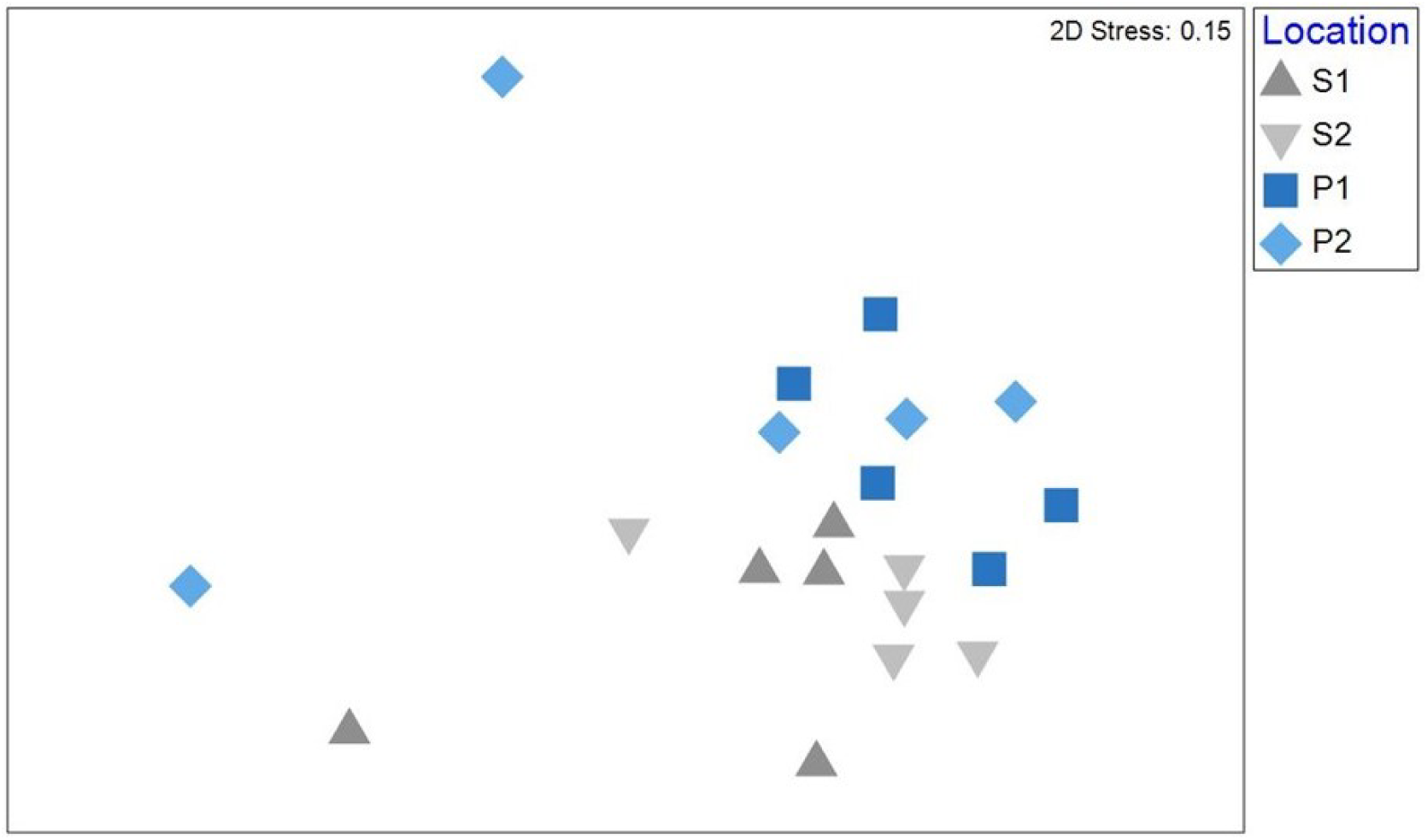
Non-metric multidimensional scaling (nMDS) plot of the chemical composition (presence/absence data, 310 features in total) of the environmental background samples per habitat type (pond/stream). The blue squares show the pond habitat type (P1, P2), while the grey triangles symbolise the stream habitat type (S1, S2).

When comparing the larval surface samples (presence/absence, excluding all features from the environmental background), we found in total 237 features that were larval-specific (figure 3). The larvae from the different habitat types shared 144 features (60,8%) and we found 54 (22.8%) of all features exclusively on stream larvae and 39 (16.5%) exclusively on pond larvae.

**Figure 3:**
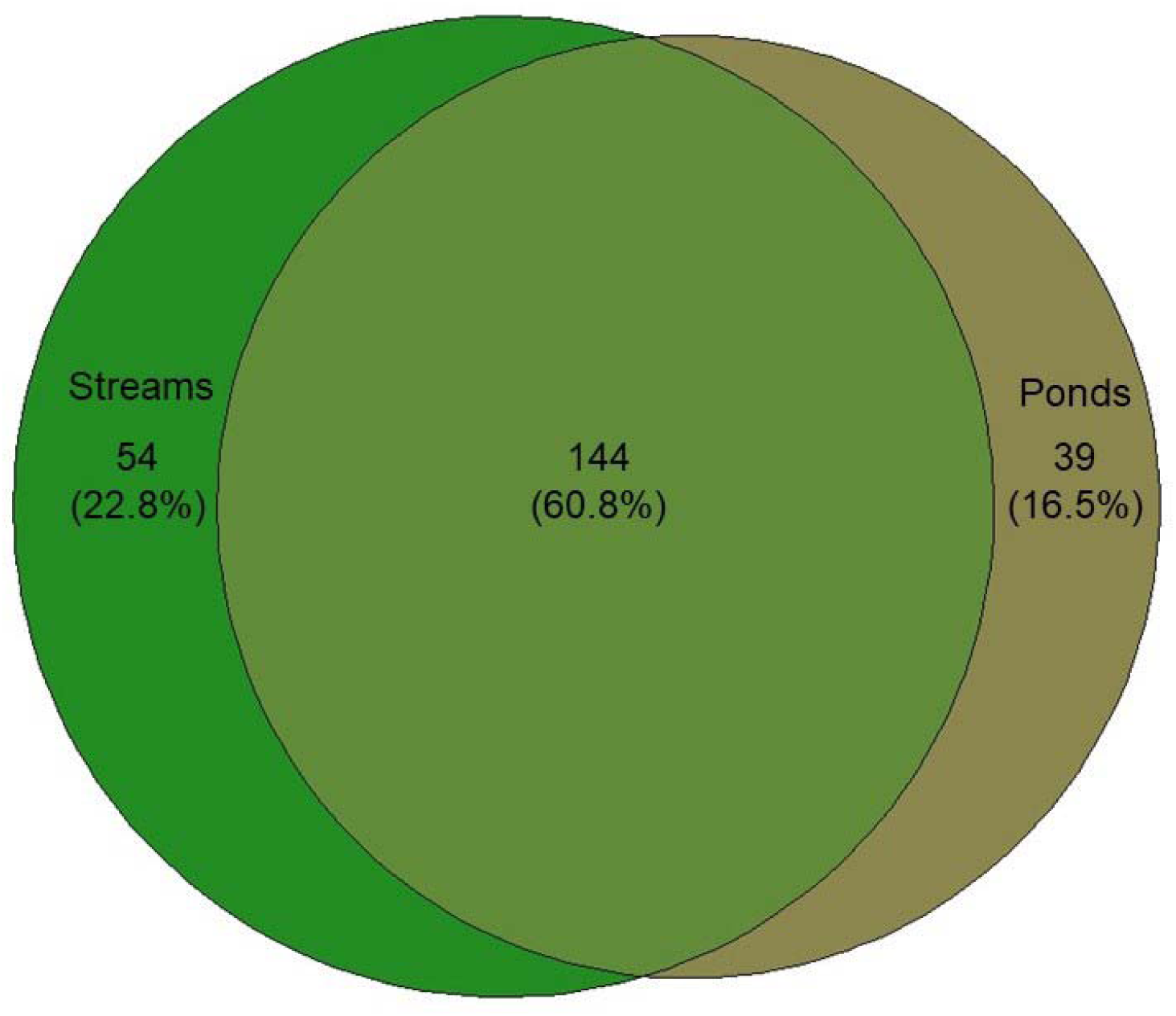
Venn diagram of the features found on the larval surface per habitat type pond or stream (based on 24 replicates per habitat type).

**Figure 4:**
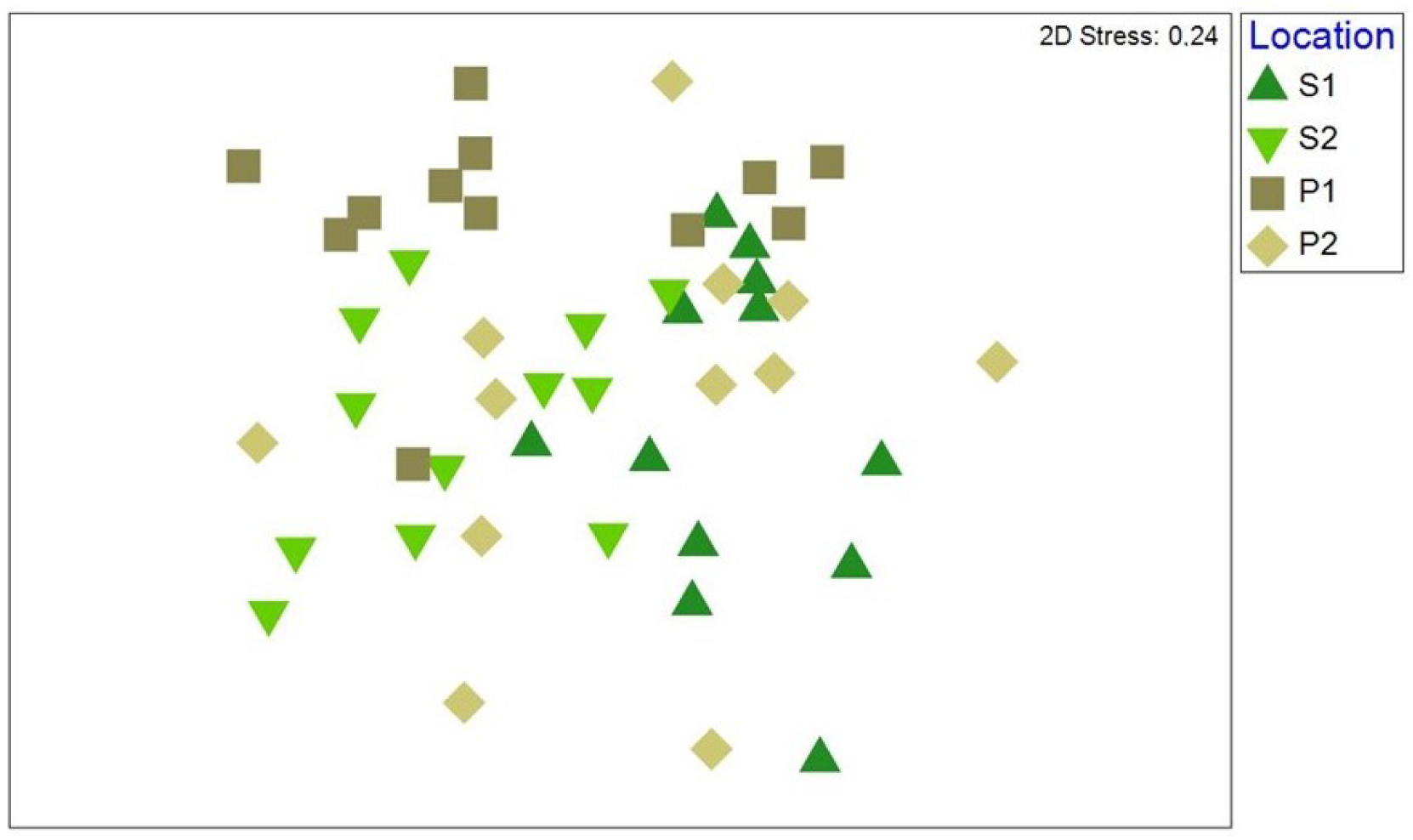
nMDS plot of the chemical composition (presence/absence data, 237 features in total) of the larval samples for each habitat type. The brown squares show the pond habitat type (P1, P2), while the green triangles symbolise the stream habitat type (S1, S2).

**Figure 5:**
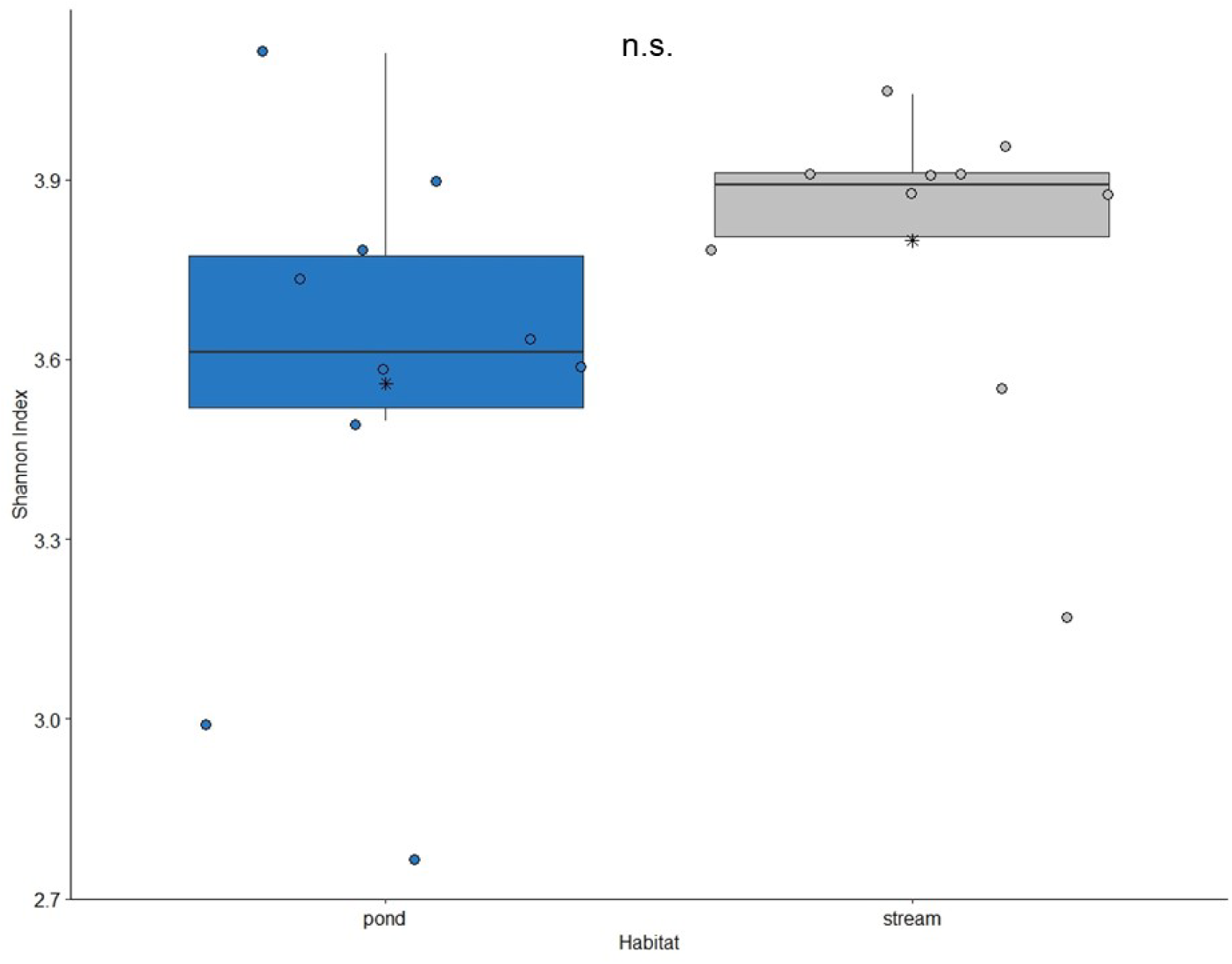
Shannon index of the features found in the environmental background samples per habitat type (pond/stream). Every point represents one sample. The horizontal line represents the median and the asterisks the mean. The boxes represent the lower and upper quartile and the whiskers the minimum and maximum values within 1.5 times the interquartile range. There was no statistically significant difference (n.s. – not significant, P > 0.05, Mann Whitney U test).

After removing all features found in the environmental background samples, we found no differences in the Shannon Index of the larval samples taken from the two habitat types (t-test: t =-0.566, p-value =0.575, Supplements figure 6).

**Figure 6:**
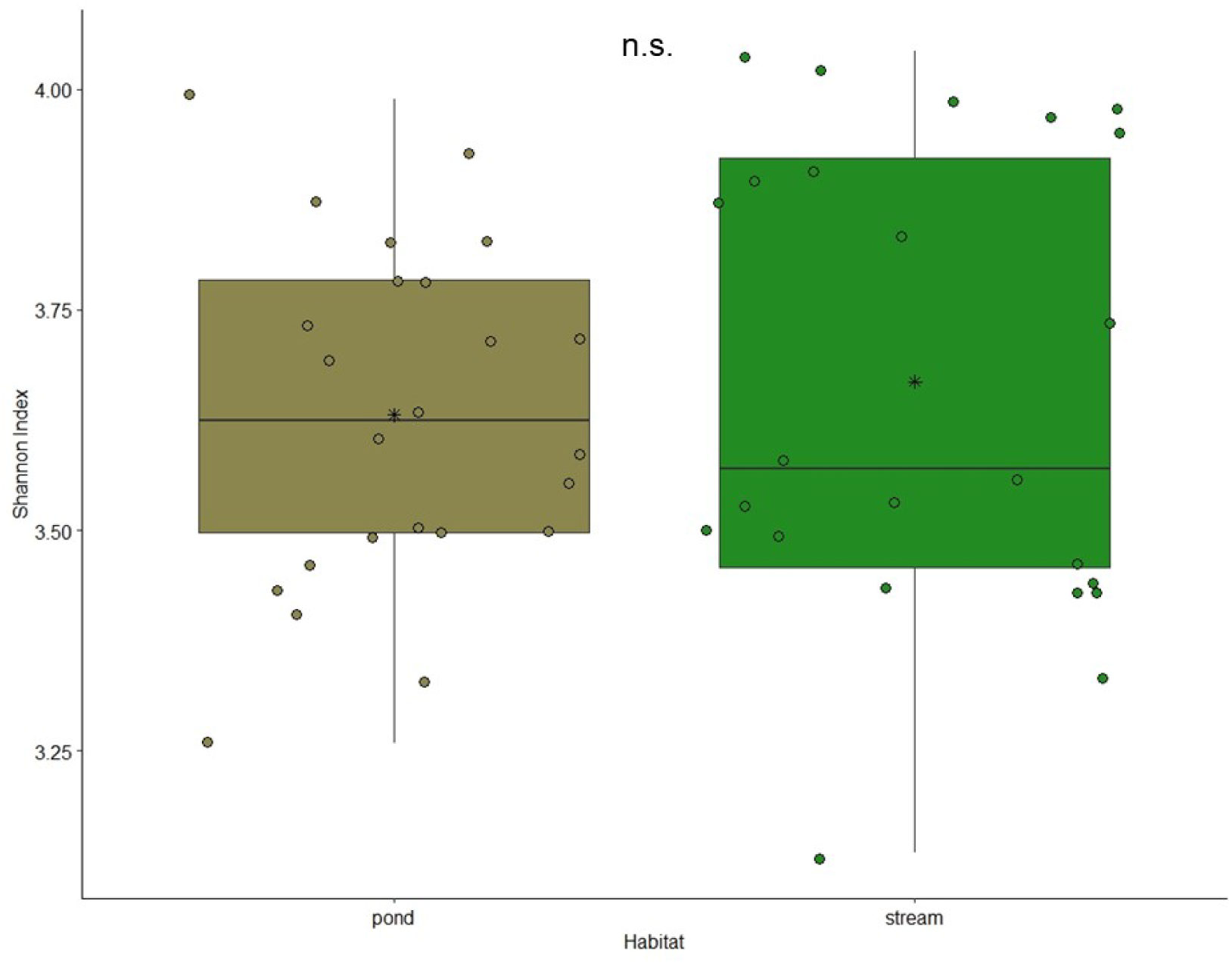
Shannon index of the features found in the larval samples per habitat type (pond/stream). Every point represents one diversity measurement. The horizontal line represents the median and the asterisks the mean. The boxes represent the lower and upper quartile and the whiskers the minimum and maximum values within 1.5 times the interquartile range. There was no statistically significant difference (n.s. – not significant, P > 0.05, t-test).

When comparing the composition of the larval samples, we found a significant difference with regard to both habitat type (one-way ANOSIM: R=0.112, p=0.002) and location (one-way ANOSIM: R=0.307, p=0.001) (figure 6).

## Discussion

We investigated potential differences in the chemical fingerprints of ponds and streams and of surfaces of fire salamander larvae collected in these habitat types. We found habitat type-specific differences when comparing the environmental background, i.e. all features that were found in the water. The presence (or absence) of these habitat type-specific features can create a distinct chemical fingerprint that can be the base for olfactory recognition. In general, habitats can have specific olfactory cues that can be perceived from larger distances by animals, such as different insect species (Webster and Cardé, 2017). For instance, the bird dwelling mosquito Culex quinquefasciatus is attracted by the smell of fresh chicken faeces, i.e. the smell of the host habitat (Cooperband et al., 2008). The aphid species Macrosiphoniella tanacetaria and Uroleucon tanaceti are able to discriminate different chemotype profiles of their host plant Tanacetum vulgare (Neuhaus-Harr et al., 2024). Many amphibian species are known to be philopatric, i.e. return as adults to the same breeding site over several years (Kusano et al., 1999). Spotted salamanders (Ambystoma maculatum), for example, are able to discriminate their pond of origin against another pond located just 3.3 km apart, using olfactory cues (McGregor and Teska, 1989).

Habitat type-specific chemical fingerprints of ponds and streams can be influenced by a variety of characteristics (Atema, 2012). Ponds and streams differ in many biotic and abiotic factors. Ponds for example often have leaf litter and terrestrial detritus on the bottom leading to constant decomposition (Reinhardt et al., 2013) which is not or in lower amounts found in streams due to the current. This decomposition could also influence the chemical features in the environmental background. Dupuis (2022) suggested that the smell of water can be influenced by pollutants, while Dacey and Wakeham (1986) describe that grazing on phytoplankton through zooplankton creates a scent environment, which is in turn used by seabirds for foraging (Nevitt et al., 1995). In addition, pH is also known to influence the survival of bacteria (Liu et al., 2023) which in turn might also influence the smell of the habitat. Similarly, the dissolved oxygen in a water body can impact bacteria. Spietz et al. (2015) found a negative correlation between dissolved oxygen and the bacterial community which might again contribute to the different chemical fingerprints in ponds and streams as they differ in their dissolved oxygen.

Our sampling design, with two locations per habitat type, allows us to draw the conclusion that ponds and streams in general differ in their chemical features. The difference between the locations emphasise this statement as the chemical composition in the different locations cluster per habitat type. The habitat type-specific characteristics might in turn directly or indirectly influence the larvae found in these habitat types. We did not aim to identify the numerous features, as we do not know which of them have a signalling function and as many of those may not be described. Furthermore, we cannot determine which of the features or which combination of them in particular are important to shape the chemical fingerprint of the larvae.

As expected, we found habitat type-specific differences in the chemical fingerprints of the surfaces of the pond and stream larvae after removing all features that we found in the environmental background. Again, we also found a location-specific effect whereby the larval surfaces from the two locations per habitat type were more similar to each other than to the larval surfaces from the location of the opposite habitat type, underlying the habitat type-specific difference. Our findings indicate that the chemical fingerprint of the larval surface is not (only) resembled by the features found in the environmental background samples. It is specific skin microbes that produce a distinct habitat type-specific smell as already known from fire salamander larvae (Bletz et al., 2016), or an adaptation to the specific habitat. Important for olfactory cues in aquatic systems are also currents, as they are often responsible to carry and disperse odour (Atema, 2012). Although fire salamander adults do not chose their larval deposition habitat based on the presence or absence of currents in the water (Krause and Caspers, 2015), it might be possible that a habitat type-specific smell is used to find the preferred larval deposition habitat type.

Moreover, the composition and abundance of macrozoobenthos differs between ponds and streams as shown by stomach content analysis of fire salamander larvae. Bletz et al. (2016) described a much greater diversity of macrozoobenthic species in streams and a very reduced diversity in ponds. Additionally, Weitere et al. (2004) found the mean energetic value of the food organisms to be much higher in streams than in ponds. Diet can influence the ability to produce signals and a good quality diet can lead to a better signal reliability (Henneken et al., 2017; Liedo et al., 2013). Thus, the availability of distinct prey items in different water bodies has the potential to influence the chemical communication of the fire salamander larvae and shape their surface “signature”.

Although our study does not allow us to draw any conclusion about the potential mechanisms involved, our findings allow speculation that the habitat type-specific features found in larval surfaces might create a habitat type-specific chemical odour in the adults. Such a habitat type-specific chemical fingerprint might be used for mate choice during assortative mating in the fire salamander (Caspers and Steinfartz, 2011), which makes the larval habitat type a potential driver for mate recognition cues. During the breeding season, the olfactory organs of adult fire salamanders are most developed, indicating the importance of olfaction during this time (Różański and Żuwała, 2020). Furthermore, the use of pheromones for mate choice and also to coordinate the male and female sexual interaction has been described from other salamander species (Woodley and Staub, 2021).

In conclusion, we found that fire salamander larvae carry habitat type-specific chemical features on their surfaces. These chemical features might be transferred from the aquatic larval habitat to the terrestrial post-metamorphic habitat and might allow habitat type-specific assortative mate recognition based on chemical cues. To test this further, we suggest a reciprocal transplant experiment of larvae between the two habitat types followed by mate choice investigated whether the smell of the habitat type is used for female navigation for larval deposition.

## Data availability

Data and code can be found https://github.com/LauraSchulte/Chemical_fingerprint.git.

## Author Contribution

BAC, CM, LF & JF designed the study; LS took the samples in the field; JF analysed the chemical composition of all samples, with input from CM; LS analysed the data and wrote the first draft of the manuscript. All authors contributed to the final manuscript.

## Conflict of Interest

We declare no conflict of interest.

## Acknowledgement

We thank Lisa Johanna Tewes and Pia Oswald for their help with the study design. We thank Eva Rousselle for her help in the field. We also thank the Behavioural Ecology Group from Bielefeld University for valuable feedback on the results.

## Funding

This research was funded by the German Research Foundation (DFG) as part of the CRC TRR 212 (NC³) – Project numbers 316099922, 396782989 and 396777092. Permissions were granted by the State Agency for Nature, Environment and Consumer Protection (LANUV; reference number: 81-02.04.2021.A437), the nature reserve authority of the Stadt Bonn and the forest warden’s office in Bonn. Experiments comply with the current laws of Germany. After the experiment, all larvae were released unharmed.

## Notes

https://github.com/LauraSchulte/Chemical_fingerprint.git

### Competing Interest Statement

The authors have declared no competing interest.

